## Supplementary Figures for "Mena regulates the LINC complex to control actin–nuclear lamina associations, trans-nuclear membrane signalling and cancer gene expression"

### Supplementary Information

Supplementary Figure 1

Supplementary Figure 2

Supplementary Figure 3

Supplementary Table 1

Supplementary Table 2

Supplementary Table 3

Supplementary Table 4

[1]Cancer Research UK Edinburgh Centre, Institute of Genetics and Cancer, University of Edinburgh, Edinburgh EH4 2XR, UK. [2]MRC Human Genetics Unit, Institute of Genetics and Cancer, University of Edinburgh, Edinburgh EH4 2XU, UK. [3]Advanced Imaging Resource, Institute of Genetics and Cancer, University of Edinburgh, Edinburgh EH4 2XU, UK. [4]Simons Initiative for the Developing Brain, School of Informatics, University of Edinburgh, Edinburgh EH8 9YL, UK. [5]Randall Centre for Cell and Molecular Biophysics, King's College London, London SE1 1UL, UK. [6]Division of Molecular and Clinical Medicine, School of Medicine, University of Dundee, Dundee DD1 4HN, UK. [7]Institute of Dentistry, Barts and the London School of Medicine and Dentistry, Queen Mary University of London, London E1 2AT, UK. *

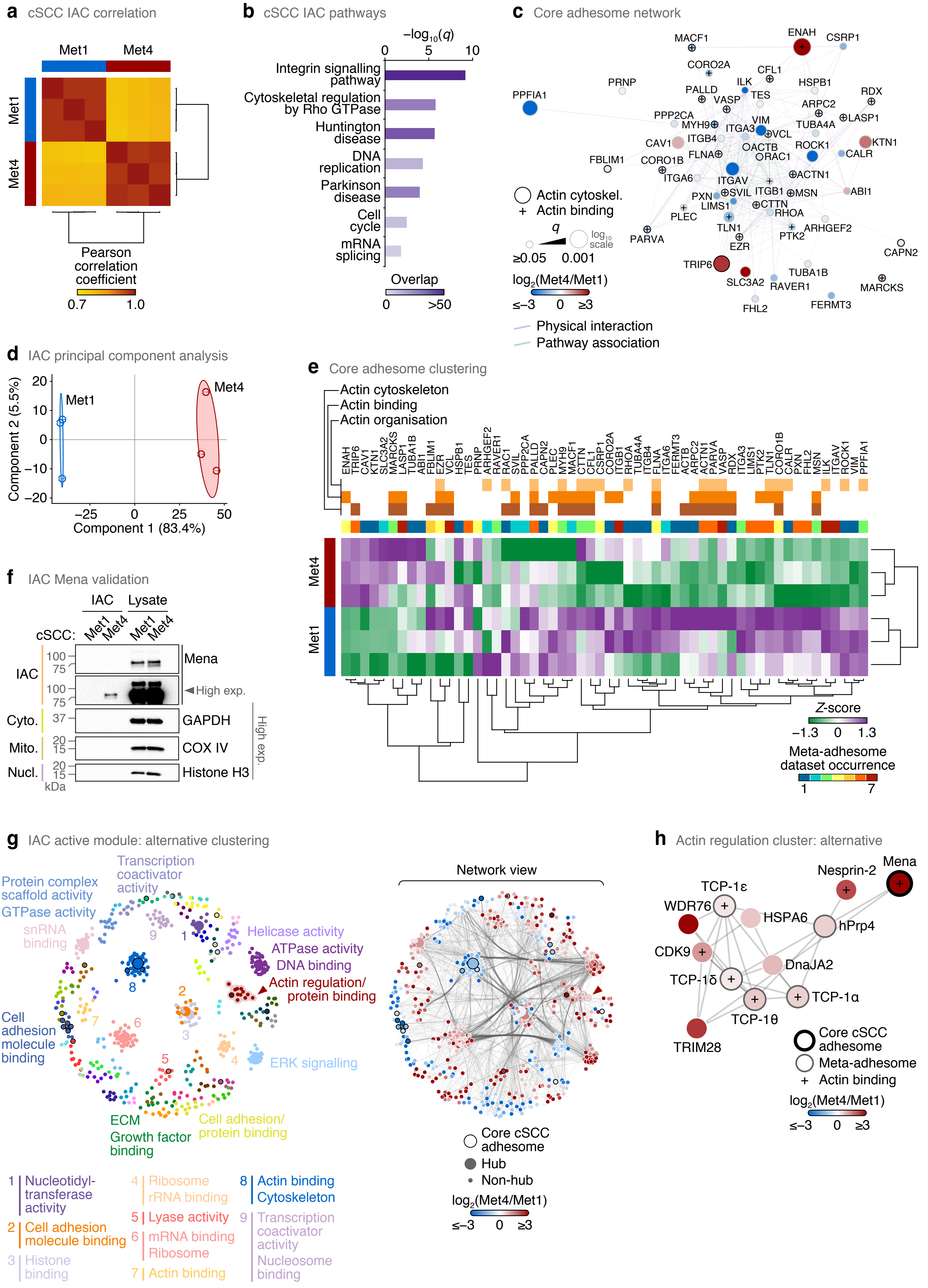


**Supplementary Figure 1.** See next page for caption.

**Supplementary Figure 1.** Proteomic analysis of patient-derived cSCC IACs. (**a**) Sample correlation analysis of mass spectrometric analyses of Met1 and Met4 IACs ($n$ = 3 independent biological replicates). Pearson correlation coefficients for all pairwise sample comparisons were subjected to hierarchical clustering. (**b**) Over-representation analysis of Panther pathways in the cSCC IAC subproteome. Purple shading intensity indicates size of subproteome overlap with respective gene sets ($q < 0.05$, hypergeometric test with Benjamini–Hochberg correction). (**c**) Functional association network of the core cSCC adhesome. Proteins (nodes) are coloured according to protein enrichment in Met1 or Met4 IACs and annotated with gene names for clarity. Node size represents significance of differential enrichment between Met1 and Met4 IACs (two-sided $t$-test with Benjamini–Hochberg correction; $n$ = 3 independent biological replicates). Black node borders indicate actin-cytoskeletal proteins; actin-binding proteins are indicated by a plus sign. (**d**) Principal component analysis of Met1 and Met4 IACs. (**e**) Hierarchical cluster analysis of the core cSCC adhesome. Data are as for Fig. 1e with additional annotation. Frequency of occurrence in meta-adhesome datasets is indicated, and proteins are annotated with gene names for clarity. (**f**) Isolation of Mena in Met4 IACs as determined by western blotting. GAPDH (cyto., cytoplasmic), COX IV (mito., mitochondrial) and histone H3 (nucl., nuclear) were probed as non-adhesion control proteins. High exp., high exposure of blot. (**g**) Maximal-scoring active module of the cSCC IAC interactome. The network is based on that in Fig. 1g, but it was partitioned using the constant Potts model, resolved for maximum Surprise quality function. Proteins (nodes) are coloured according to assigned cluster (left panel). Molecular functions enriched in protein clusters were determined by Gene Ontology over-representation analysis (the two most significant functional categories are labelled). For protein clusters with fewer than two significant functional categories, clusters were annotated manually with a single representative term. Network view (right panel) shows corresponding protein interactions (edge densities) in the partitioned network; nodes are coloured according to protein enrichment in Met1 or Met4 IACs. The actin regulation cluster detailed in **h** is indicated with a red arrowhead. Large nodes indicate kinless and connector hubs; black node borders indicate core adhesome proteins. (**h**) Subnetwork analysis of the actin regulation cluster identified by active module partitioning in **g**. The weighted subnetwork was clustered using a force-directed algorithm. Nodes are coloured according to protein enrichment in Met1 or Met4 IACs.

---

**a** Mena re-expression

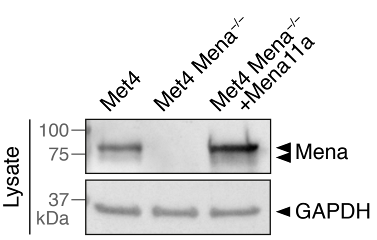


**b** Mena sequence alignment

```
Met4      GGATCAACAATAGAAACAGAACAAAAAGAGGACAAAGGTGAAGATTCAGAGCCTGTAACT  355
Pan-Mena  GGATCAACAATAGAAACAGAACAAAAAGAGGACAAAGGTGAAGATTCAGAGCCTGTAACT  1500
Mena11a   GGATCAACAATAGAAACAGAACAAAAAGAGGACAAAGGTGAAGATTCAGAGCCTGTAACT  1500
          ************************************************************

Met4      TCTAAGGCCTCTTCAACAAGTACACCTGAACCAACAAGAAAACCTTGGGAAAGAACAAAT  415
Pan-Mena  TCTAAGGCCTCTTCAACAAGTACACCTGAACCAACAAGAAAACCTTGGGAAAGAACAAAT  1560
Mena11a   TCTAAGGCCTCTTCAACAAGTACACCTGAACCAACAAGAAAACCTTGGGAAAGAACAAAT  1560
          ************************************************************

Met4      ACAATGAATGGCAGCAAGTCACCTGTTATCTCCAGACGGGATTCTCCAAGGAAAAATCAG  475
Pan-Mena  ACAATGAATGGCAGCAAGTCACCTGTTATCTCCAGA------------------------  1596
Mena11a   ACAATGAATGGCAGCAAGTCACCTGTTATCTCCAGACGGGATTCTCCAAGGAAAAATCAG  1620
          ************************************

Met4      ATTGTTTTTGACAACAGGTCCTATGATTCATTACACAGACCAAAATCCACACCCTTATCA  535
Pan-Mena  ---------------------------------------CCAAAATCCACACCCTTATCA  1617
Mena11a   ATTGTTTTTGACAACAGGTCCTATGATTCATTACACAGACCAAAATCCACACCCTTATCA  1680
                                                 *********************

Met4      CAGCCCAGTGCCAATGGAGTCCAGACGGAAGGACTTGACTATGACAGGCTGAAGCAGGAC  595
Pan-Mena  CAGCCCAGTGCCAATGGAGTCCAGACGGAAGGACTTGACTATGACAGGCTGAAGCAGGAC  1677
Mena11a   CAGCCCAGTGCCAATGGAGTCCAGACGGAAGGACTTGACTATGACAGGCTGAAGCAGGAC  1740
          ************************************************************

Met4      ATTTTAGATGAAATGAGAAAAGAATTAACAAAGCTAAAAGAAGAGCTCATTGATGCAATC  655
Pan-Mena  ATTTTAGATGAAATGAGAAAAGAATTAACAAAGCTAAAAGAAGAGCTCATTGATGCAATC  1737
Mena11a   ATTTTAGATGAAATGAGAAAAGAATTAACAAAGCTAAAAGAAGAGCTCATTGATGCAATC  1800
          ************************************************************
```

**Supplementary Figure 2.** Analysis of Mena expression in Met4 cSCC cells. (**a**) Western blot analysis of endogenous Mena in parental Met4 cells, absence of Mena in Mena-depleted Met4 cells (Met4 Mena$^{-/-}$) and re-expression of Mena11a in Met4 Mena$^{-/-}$ cells (Met4 Mena$^{-/-}$ +Mena11a). (**b**) Nucleotide sequence alignment of endogenous Mena transcript sequenced from Met4 cells. Mena sequence compared to pan-Mena (GenBank accession KM214203.1) and Mena11a (GenBank accession KM214204.1) coding sequences. Grey boxes indicate the additional Mena11a exon coding sequence. Asterisks indicate exact residue conservation.

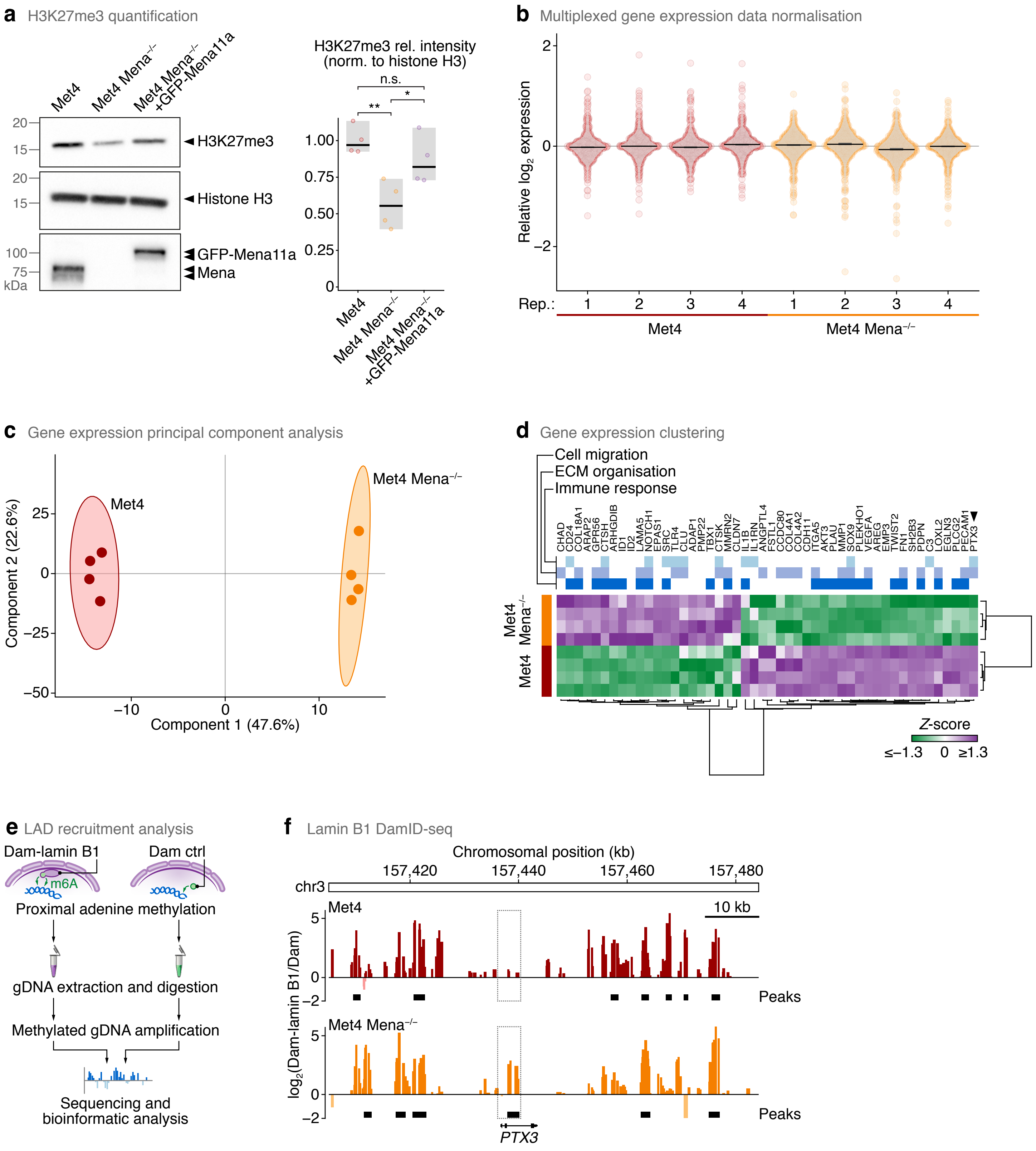


**Supplementary Figure 3.** See next page for caption.

**Supplementary Figure 3.** Regulation of cSCC gene expression by Mena. (**a**) Western blot analysis of H3K27me3 in Met4, Met4 Mena$^{-/-}$ and Met4 Mena$^{-/-}$ +GFP-Mena11a cell lysates. Densitometric intensities were normalised (norm.) to histone H3 and expressed relative (rel.) to Met4 cells (right panel). Black bar, median; light grey box, range. **$P < 0.01$, *$P < 0.05$; n.s., not significant; one-way ANOVA with Tukey's correction ($n = 4$ independent experiments). (**b**) Normalised cancer progression gene expression data across independent biological replicate (rep.) analyses for Met4 and Met4 Mena$^{-/-}$ cells. Black bar, median; dark grey box, 95% confidence interval; light grey silhouette, probability density. Most 95% confidence intervals are smaller than the weight of the corresponding median line (black bar) and so are not visible. (**c**) Principal component analysis of cancer progression gene expression for Met4 and Met4 Mena$^{-/-}$ cells. (**d**) Hierarchical cluster analysis of cancer progression genes significantly differentially regulated between Met4 and Met4 Mena$^{-/-}$ cells ($q < 0.05$, two-sided $t$-test with Benjamini–Hochberg correction; $n = 4$ independent biological replicates). Data are as for Fig. 4d with additional annotation. *PTX3* is indicated with a black arrowhead. (**e**) Workflow for identification of recruitment of lamina-associated domains (LADs) of chromatin to the nuclear lamina by DamID. DNA adenine methyltransferase (Dam) tethered to lamin B1 or untethered Dam control (ctrl) was expressed in Met4 and Met4 Mena$^{-/-}$ cells. Dark blue strands, genomic DNA (gDNA); green arrows, adenine methylation (m6A) in GATC motifs proximal to Dam. (**f**) Nuclear lamina association profiles revealed by lamin B1 DamID sequencing (DamID-seq). Lamin B1 DamID-seq tracks were generated from Met4 (red, top profile) and Met4 Mena$^{-/-}$ (orange, bottom profile) cells, and an 80-kb region of chromosome 3 (human reference genome GRCh38) spanning the *PTX3* gene is shown. The highest-scoring putative enhancer region associated with *PTX3* (GeneHancer identifier GH03J157436) is indicated with a dotted box. Significantly called DamID peaks are indicated with black bars (FDR < 5%). Scale bar, 10 kb.
